# The microtubule-associated proteins CKAP2 and its paralog CKAP2- Like control ciliogenesis in human cells

**DOI:** 10.64898/2026.09.03.749104

**Authors:** Luiz E. da Silva, Gabriel Campolina-Silva, Jenine Afani, Clémence Belleannée, Susanne Bechstedt

**Affiliations:** Department of Anatomy and Cell Biology and Centre de recherche en Biologie Structurale, McGill University, Montreal, Canada; CHU de Québec Research Center (CHUL), Université Laval, Quebec City, Canada; Centre de recherche en Reproduction, Développement et Santé Intergénérationnelle, Université Laval, Quebec City, Canada

**Keywords:** Primary cilia, Microtubules, Microtubule-associated proteins, Paralogous compensation

## Abstract

Ciliogenesis is an evolutionarily conserved process that leads to the assembly of cilia. This process relies on microtubule-associated proteins (MAPs) to regulate axonemal microtubule dynamics. Misregulation of MAPs often leads to changes in ciliary homeostasis, contributing to numerous ciliopathies. Although CKAP2 and CKAP2-Like are best known for regulating microtubule dynamics during cell division, they are also components of ciliary organelles. Notably, CKAP2 overexpression promotes chromosomal instability and cancer, whereas CKAP2-Like deficiency causes Filippi syndrome, a developmental disorder with clinical features overlapping those of ciliopathies. How both MAPs operate at primary cilia remains poorly understood. Here, we identify both MAPs as axonemal components of motile and primary cilia in human cells. We find that CKAP2-positive cilia are linked to cell cycle progression and that the conserved intrinsically disordered microtubule-binding domain of CKAP2, which is essential for microtubule polymerization and stabilization, is required for its ciliary localization. Unexpectedly, CRISPR-mediated loss of either paralog results in elongated cilia without affecting ciliogenesis, accompanied by increased ciliary enrichment of the remaining paralog and impaired Gli2 accumulation at the ciliary tip. In contrast, simultaneous deletion of both MAPs suppresses ciliogenesis and has minimal effect on the ciliary length as compared to wild-type levels. These findings uncover that CKAP2 and CKAP2-Like are axonemal MAPs that contribute to ciliogenesis, partially compensating for each other in human cells.

## Introduction

Ciliogenesis is an evolutionarily conserved process by which cells assemble cilia. Cilia are microscopic, hair-like projections that extend from the cell surface and perform several essential functions. They act by propelling cells, sweeping fluids or particles across surfaces and by converting extracellular cues into intracellular signaling events. There are two types of cilia, motile and non-motile, which differ in structure and function (Mill et al., 2023; Satir and Christensen, 2007). Primary cilia are often referred to as non-motile, sensory organelles responsible for regulating many signaling pathways during embryonic development and in differentiated tissues (Anvarian et al., 2019). To properly perform their function, primary cilia have a backbone formed by a microtubule-based structure known as the axoneme, which is anchored by the basal body and surrounded by a specialized membrane (Mill et al., 2023). In mammalian cells, this axoneme consists of nine outer microtubule doublets in the proximal region, which transition into singlets toward the middle-distal end (Kiesel et al., 2020). Axoneme growth and stabilization are supported by microtubule-associated proteins (MAPs) (Zhang et al., 2025), microtubule inner proteins (MIPs) (Gui and Orbach, 2025) and post-translational modifications of tubulin (Wloga et al., 2017). Interestingly, microtubule plus-ends at primary cilia exhibit slow turnover rates, which have recently been recapitulated by a set of ciliary tip interacting MAPs (Saunders et al., 2025). Recent progress in the cilia field has focused on defining the structural basis of axonemes(Leung et al., 2025; Ma et al., 2019) and in understanding the intrinsic heterogeneity of primary cilia across different cell types and even within the same population (Hansen et al., 2025). Despite that, considerably less is known about MAPs that act dynamically along the axoneme, or how related MAPs may provide functional robustness to this compartment. Notably, many of them are also components of the mitotic spindle, thereby connecting primary cilia dynamics to cell cycle progression and division (Doornbos and Roepman, 2021).

CKAP2 and its paralog CKAP2-Like were first identified as MAPs involved in the regulation of the mitotic spindle (Maouche-Chrétien et al., 1998; Yumoto et al., 2013). Our previous research has demonstrated that CKAP2 controls microtubule growth rate *in vitro* (McAlear and Bechstedt, 2022) and in cells (Paim et al., 2024). Its misregulation leads to error-prone chromosomal segregation, contributing to cancer development (Jeon et al., 2006; Jin et al., 2004; Paim et al., 2024; Tsuchihara et al., 2005; Yoo et al., 2016). More recently, we have elucidated the underlying mechanisms by which CKAP2 promotes microtubule growth and stabilization, proposing that it functions as a polymerase (Lyalina et al., 2025). On the other hand, loss-of-function mutations in CKAP2-Like cause Filippi syndrome (Hussain et al., 2014), a rare developmental disorder characterized by growth defects, intellectual disability, syndactyly and microcephaly (Hussain et al., 2014). Due to the similarities in clinical manifestations with ciliopathies commonly associated with defects in primary cilia (Deretic et al., 2023), we hypothesize that CKAP2-Like may also contribute to Filippi syndrome by regulating primary cilia. Both MAPs have previously been identified as candidate ciliary-associated proteins through unbiased proteomic and functional screening datasets (Gupta et al., 2015; Mick et al., 2015). In line with this, CKAP2-Like has been recently detected in the primary cilia of the developing mouse brain (Liu et al., 2026). Its knockout results in microcephaly, impaired Hedgehog signaling (Liu et al., 2026) and male infertility issues in mice (Lyu et al., 2025). How CKAP2 and CKAP2-Like contribute to ciliary homeostasis, whether individually or in combination, remains poorly understood.

Here, we analyzed the presence of both MAPs in the ciliated cells of the human male reproductive tissue. We found that they are distributed along the axonemes and basal bodies of the motile and primary cilia of the human efferent ductules and epididymis, suggesting an association with specialized microtubules in a conserved pattern. We further analyzed CKAP2 and CKAP2-Like’s presence in RPE-1 cells and found differences in their localization pattern, which correlate with the cell cycle stage. By combining CRISPR/Cas9-genome editing and live cell imaging, we captured the dynamics of CKAP2 localization following primary cilia assembly and throughout their disassembly prior to mitosis. We also demonstrated that cells lacking CKAP2 or CKAP2-Like develop elongated cilia due to mutual compensation, while concomitant depletion of both MAPs negatively impacts ciliogenesis. Our findings further establish that CKAP2 and CKAP2- Like are important regulators of axoneme assembly and maintenance, likely having overlapping functions in human cells.

## Results

### CKAP2 and CKAP2-Like localize to motile and primary cilia in the human efferent ductules and epididymis

We and others have previously observed consistent ciliary staining of CKAP2 in mammalian cells (Liu et al., 2026; Lyu et al., 2025; Paim et al., 2024), which prompted us to explore its localization and potential role at primary and motile cilia. The human male reproductive tract contains cells with both primary and motile cilia, in addition to sperm flagella. In this system, efferent ductules are small tubes that concentrate and carry sperm from the testes to the epididymis, where sperm cells become functionally mature (Augière et al., 2024; Vinay et al., 2025). We thus probed for the presence of both MAPs in this system, since it has been recently shown that CKAP2-Like mRNA and protein levels are enriched in the male reproductive tract of mice (Lyu et al., 2025). Notably, we found that CKAP2 and CKAP2-Like showed similar staining patterns in the human efferent ductules and epididymis (Fig. 1*A* and *B*, respectively; Fig. S1*A* and *B*). In the efferent ductule epithelium, multiciliated cells (MMCs) were readily identified by the strong acetylated α- tubulin signal along the axonemes of their numerous motile cilia, as well as by the multiple γ-tubulin-positive centrioles forming the basal bodies (right inset, Fig. 1*A*; Fig. S1*A*). In these cells, both CKAP2 and CKAP2-Like showed a relatively diffuse intracellular distribution, but their staining was more prominent in the ciliary compartment. Both MAPs were detected at the basal bodies and along the ciliary axonemes, occasionally localizing to the ciliary tip (right inset, Fig. 1*A;* Fig. S1*A*).

**Figure 1.**
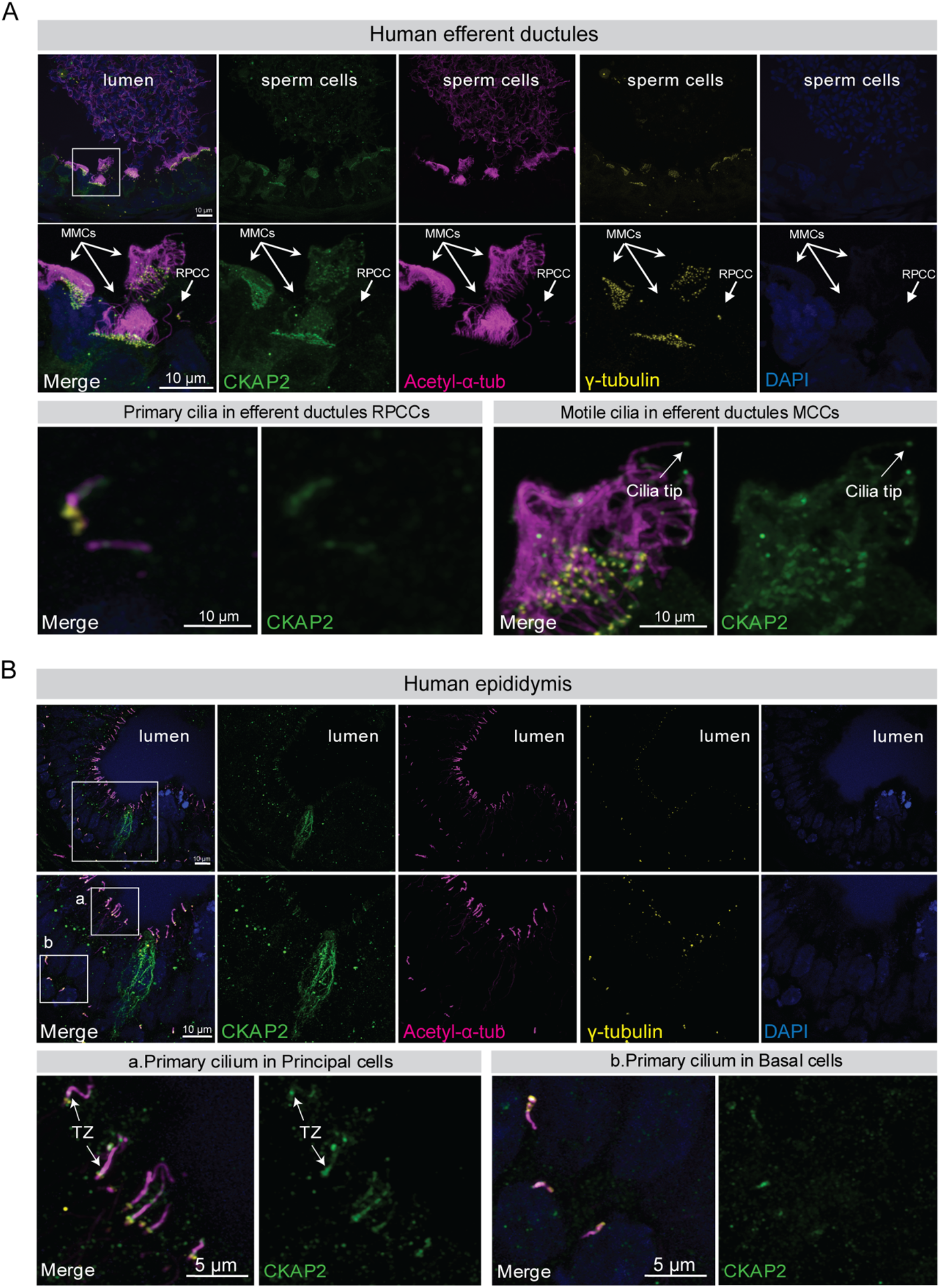
CKAP2 is found in the motile and primary cilia of the human male reproductive tract. **(A)** Confocal imaging microscopy of human efferent ductules stained for CKAP2, acetylated- α-tubulin (axoneme) and γ-tubulin (basal bodies). Insets indicate CKAP2 localization in multiciliated cells (MMCs), which are characterized by the apical alignment of numerous γ-tubulin-positive basal bodies, and in reabsorptive primary ciliated cells (RPCC). Occasional CKAP2 localization at the ciliary tip can be observed in MMCs. CKAP2 is also present in sperm flagella of the duct lumen. **(B)** Confocal imaging microscopy of human epididymis stained for CKAP2, acetylated-α-tubulin (axoneme) and γ-tubulin (basal bodies). Insets indicate CKAP2 localization in the primary cilia of principal and basal cells. Occasional CKAP2 localization at the transition zone (TZ) of primary cilia can be observed in principal cells.

We next examined the solitary primary cilia found in epithelial cells neighboring the MMCs of the efferent ductules, which correspond to reabsorptive primary ciliated cells (RPCCs), previously known as non-ciliated efferent ductules cells (Augière et al., 2024; Vinay et al., 2025). Although these solitary cilia were more difficult to distinguish from the surrounding motile cilia, both CKAP2 and CKAP2-Like were detected at the basal body and along the axoneme and ciliary membrane (left inset, Figure 1A; Fig. S1*A*). In RPCCs in which a primary cilium was not morphologically evident, both proteins were detected at the centrosome, as indicated by their colocalization with γ-tubulin.

We then examined the caput epididymis, where primary cilia could be readily identified by acetylated α-tubulin staining in both the apical epithelial compartment, associated with principal cells (a, Fig. 1*B*; Fig. S1*B*), and the basal compartment, where basal epithelial cells reside (b, Fig. 1*B*; Fig. S1*B*). Similar to the primary cilia observed in the efferent ductules, CKAP2 and CKAP2- Like were detected at the basal body and along the axoneme and ciliary membrane. In non-ciliated cells, both proteins were detected at the centrosome and colocalized with γ-tubulin (Fig. 1*B*). Interestingly, both MAPs display prominent accumulation at the transition zone (TZ), and occasionally at the ciliary tip.

Finally, CKAP2 and CKAP2-Like labeling could also be seen in the flagella of spermatozoa present in the lumen of the efferent ductules (Fig. 1*A*). However, the staining appears to be lower in intensity when compared to the staining pattern of CKAP2 and CKAP2-Like at the primary and motile cilia. Thus, CKAP2 and CKAP2-Like were detected in all three ciliary structures examined in the human male reproductive tract—motile cilia, primary cilia, and sperm flagella—with staining observed at the basal body and along the axoneme and/or ciliary membrane.

### CKAP2 localization to the primary cilia is linked to cell cycle progression

We next used immortalized human retinal pigment epithelial (hTERT RPE-1) cells, a well-established model system, to investigate CKAP2-related ciliogenesis (Ford et al., 2018; Hansen et al., 2025; Spalluto et al., 2013). To this end, cells were serum-starved for 48 h and co-stained for endogenous CKAP2 and ciliary axonemes (acetylated-α-tubulin) (Fig. 2*A*). Under this particular condition, we found that CKAP2 localized to approximately 25% of the total ciliated cells (Fig. 2*A* and *B*). Interestingly, in individual cells CKAP2 was either absent from primary cilia (a, Fig. 2*A*), decorated only the axoneme (b, Fig. 2*A*) or extended toward the basal body and the cytoplasmic microtubule network (c, Fig. 2*A*). A similar localization pattern was observed for the paralog CKAP2-Like (Fig. S2*A*). Remarkably, CKAP2-positive cells exhibited lower levels of tubulin acetylation within cilia (Fig. 2*C*) as well as shorter cilia (Fig. 2*D*) compared to CKAP2-negative cells. Since cell cycle progression is coupled to both CKAP2 expression(Paim et al., 2024) and deacetylation of primary cilia with concomitant reduction in ciliary length (Liang et al., 2016; Ran et al., 2015; Sung and Li, 2011), we reasoned that CKAP2-positive cells were further along in the cell cycle than CKAP2-negative cells. Although serum starvation is known to promote cell cycle arrest (Sung and Li, 2011), we observed a proportion of cells that continues to divide, with CKAP2 localizing to the centrosomes and mitotic spindle (d, Fig. 2*A*). To confirm the correlation between cell-cycle stage and CKAP2 localization to cilia, we co-stained cells for the proliferation marker Ki67 and CKAP2 (Fig. 2*E*). Based on this dual-staining strategy, we classified three major patterns for CKAP2 expression across the cell cycle. In quiescent cells (i.e. negative Ki67), CKAP2 was absent from primary cilia, whereas in cycling cells (i.e. positive Ki67), CKAP2 distributed either along the axoneme (45.0%) or also decorated the cytoplasmic microtubules (55.0%) (Fig. 2*E*). Amongst CKAP2-populated cilia, we observed a positive correlation between its expression and nuclear Ki67 levels, indicating that CKAP2’s presence at primary cilia becomes more prominent as cells progress through the cell cycle, with additional decoration of cytoplasmic microtubules at later stages (i.e. G2 phase) (Fig. 2*F*). Altogether, these data demonstrate that CKAP2’s dynamic localization within the primary cilia is closely related to cell cycle progression (Fig. 2*G*).

**Figure 2.**
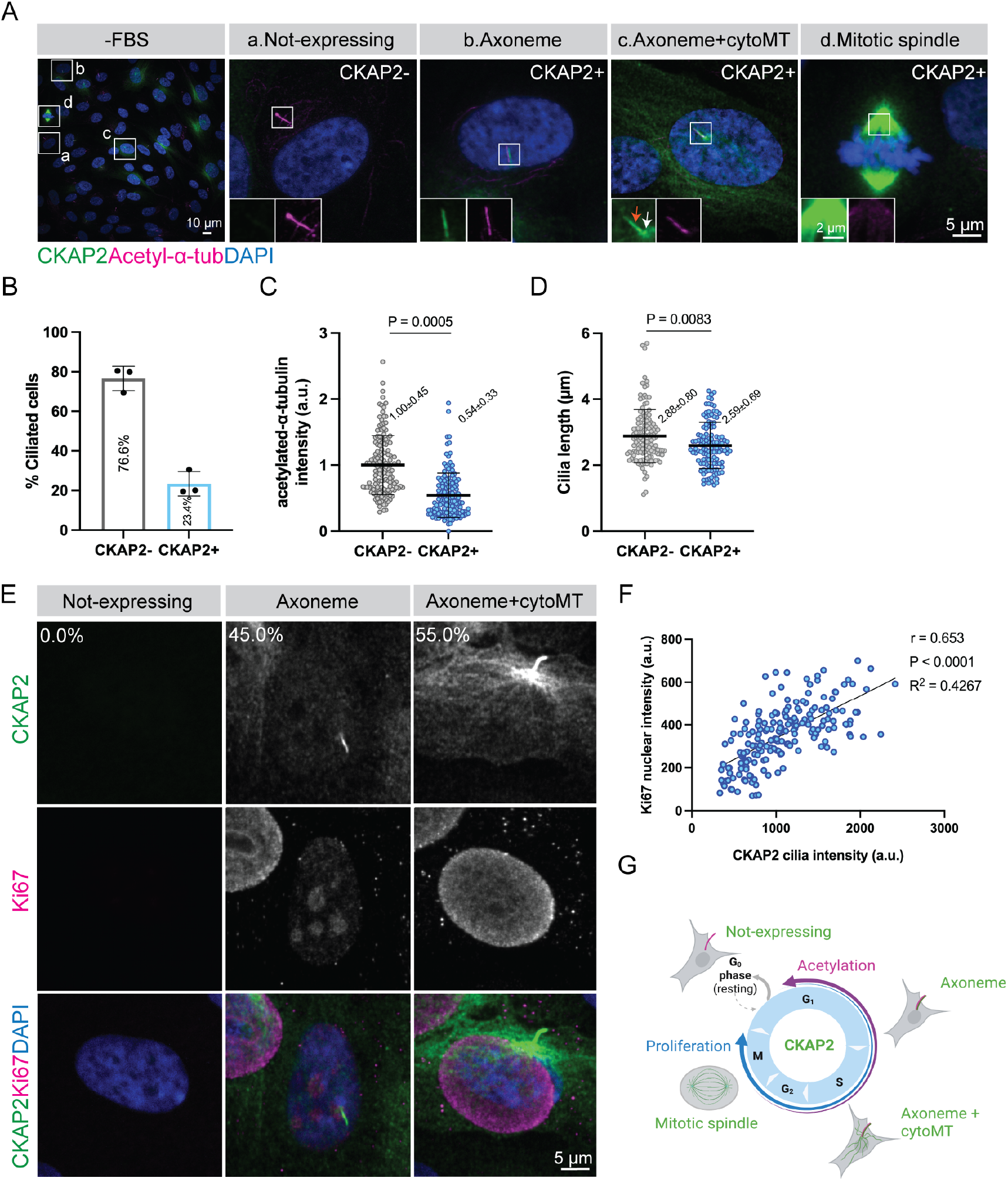
CKAP2 is found in the primary cilia of cycling RPE-1 cells. **(A)** Wild-type cells were serum starved for 48 h (-FBS), fixed and co-stained for endogenous CKAP2 and acetylated-α- tubulin. Insets indicate different CKAP2 localization patterns, including axoneme (orange arrow) and basal body (white arrow). **(B)** Quantification of the percentage of cells in which CKAP2 localizes to primary cilia as determined by the axonemal marker acetylated α-tubulin (n = 500 cells). Each dot represents an independent experiment (N = 3). **(C)** Levels of acetylated α-tubulin at primary cilia in CKAP2-positive (n = 138 cells) and CKAP2-negative cells (n = 151 cells). Results are representative of at least two independent experiments. **(D)** Ciliary length comparison between CKAP2-expressing (n = 118 cells) and non-expressing cells (n = 128 cells). Results are representative of at least two independent experiments. **(E)** Serum-starved cells were co-stained for CKAP2 and Ki67, then classified according to the different CKAP2 localization patterns observed in panel A and presented as a percentage of the total cell population (n = 271 cells). **(F)** Pearson correlation between Ki67 nuclear expression and CKAP2 expression at primary cilia (n = 193 cells). **(G)** Schematic representation of CKAP2’s dynamic localization throughout the cell cycle. P < 0.05 (statistically significant). Nuclei were stained with DAPI. Measurements are reported as average ± SD.

### CKAP2 dynamically localizes to primary cilia during their assembly and disassembly

Previous studies have shown that primary cilia persist throughout the S and G2 phases and are resorbed shortly before mitosis (Ford et al., 2018), which coincides with the timing of CKAP2 expression reported here (Fig. 2*E-G*) and elsewhere (Paim et al., 2024). To gain further insight into the spatiotemporal distribution of CKAP2 within primary cilia, we performed genome editing using CRISPR-Cas9 knock-in to introduce an N-terminal Green Fluorescent Protein (GFP)- tag to *CKAP2* in RPE-1 cells (Paim et al., 2024). We then used CenSpark-650, a recently developed ciliary probe, to label the microtubule doublets and triplets within the centrioles and basal bodies of these cells, as well as the proximal region of their cilia (Pourroy et al., 2026). Notably, the CenSpark signal decreases toward the ciliary tip, likely due to the shift to singlet microtubules in that region (Kiesel et al., 2020; Pourroy et al., 2026). We found that the endogenous CKAP2 signal peaked distal to the CenSpark probe and then declined in proportion to singlet microtubule density (Kiesel et al., 2020) (Fig. S2*B* and *C*). Our data suggest that CKAP2 may predominantly act on singlet microtubules in primary cilia rather than the doublet axoneme microtubules.

Next, we conducted extensive live imaging with CenSpark-650 to monitor ciliary dynamics prior to mitotic entry. After the initial stages of axoneme assembly, we observed that CKAP2 reaches the ciliary tip, where it becomes incorporated and gradually extends towards the ciliary base (Fig. 3*A* and Movie S1). Using the same live-imaging approach, we further tracked CKAP2 localization during ciliary disassembly. This process involves making a complex decision between rapid deciliation, gradual resorption or a combination of both (Mirvis et al., 2019). We found that CKAP2 follows the depolymerization of the axoneme prior to mitosis (Fig. 3*B* and Movie S2). It then accumulates in duplicated centrosomes and populates the microtubules of the mitotic spindle during metaphase, before shifting to the chromatin at mid-anaphase and being degraded at the end of mitosis. Overall, these experiments provide an overview of CKAP2 localization in relation to ciliary dynamics throughout the cell cycle.

**Figure 3.**
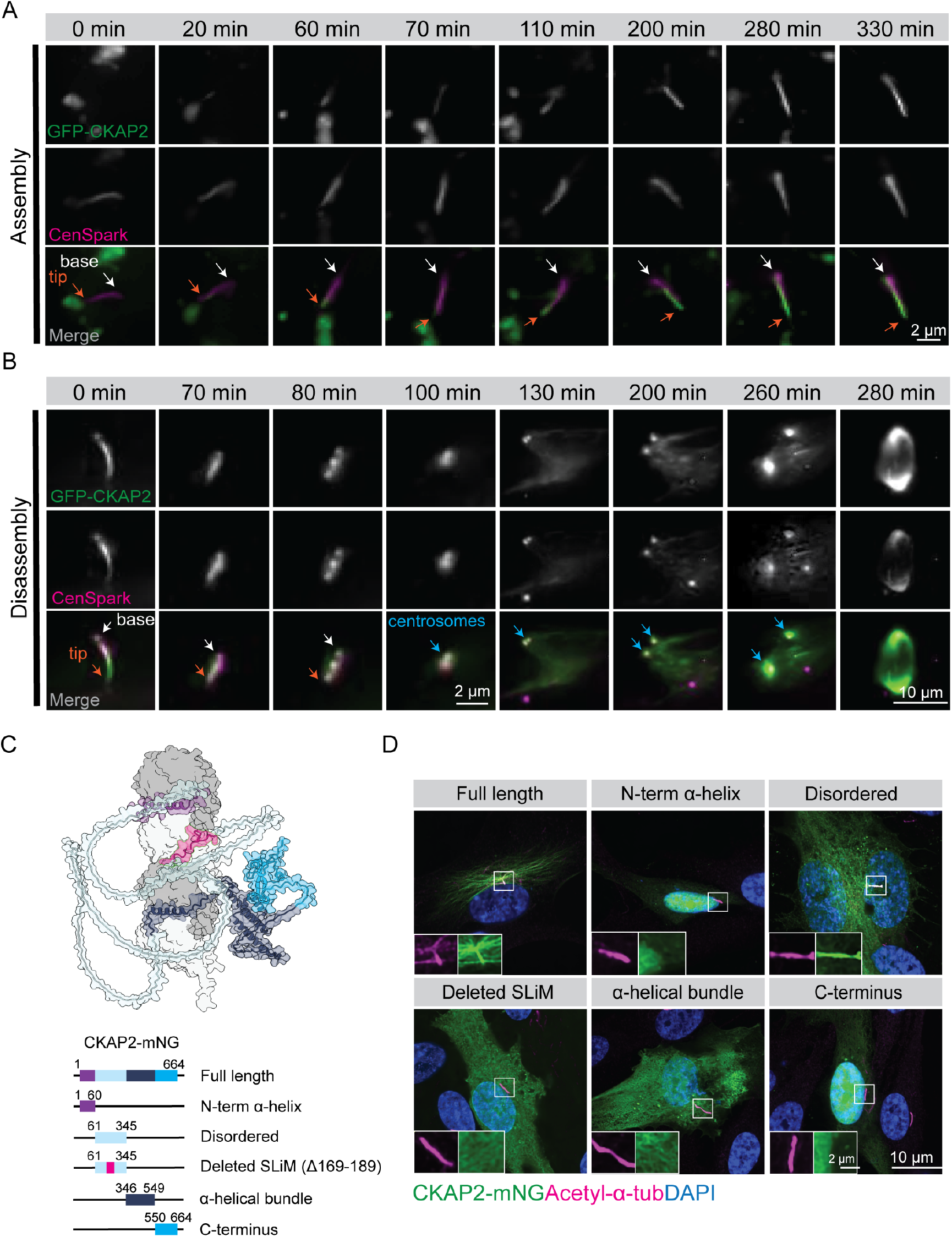
Dynamics of CKAP2 localization to primary cilia in cycling cells. **(A)** Excerpt of time-lapse imaging showing the dynamic localization of CKAP2 within primary cilia following their assembly. GFP-CKAP2 knock-in RPE-1 cells were serum starved for 32 h and imaged every 10 min for 20 h (see Movie S1). The ciliary probe CenSpark-650 was added to media to visualize the basal bodies and axoneme. **(B)** Excerpt of time-lapse imaging showing the dynamic localization of CKAP2 within primary cilia during their disassembly. GFP-CKAP2 knock-in RPE-1 cells were serum starved for 32 h and imaged every 10 min for 20 h (see Movie S2). Orange and white arrows indicate the ciliary tip and base, respectively. Blue arrows indicate the centrosomes. **(C)** Schematic representation of CKAP2 constructs expressed in RPE-1 cells (lower) based on AlphaFold3 model of CKAP2 and two tubulin dimers (upper). **(D)** Cells were transiently transfected with CKAP2 (full length) or CKAP2 fragments for 24 h, serum-starved for 48 h, fixed and stained for acetylated-α- tubulin. Nuclei were stained with DAPI. Data are representative of at least two independent experiments.

### The disordered microtubule-binding domain of CKAP2 is required for its localization to primary cilia

We further sought to determine which domains of CKAP2 are required for its localization to primary cilia. Modelling predictions of CKAP2 and two tubulin dimers generated by AlphaFold3 suggested four distinct domains within CKAP2 structure: the N-terminal α-helix, the disordered domain, the central α-helical bundle and the C-terminal domain. We have recently demonstrated that the disordered domain of CKAP2 harbors a highly conserved Short Linear Motif (SLiM), which is responsible for its microtubule-binding and stabilizing activity (Lyalina et al., 2025). We thus transiently expressed constructs encoding these four domains in RPE-1 cells (Fig. 3*C*). In addition to CKAP2-full length, cells expressing only the disordered domain of CKAP2 retained ciliary localization (Fig. 3*D*). Conversely, a construct that lacks the 21-amino acid SLiM within CKAP2’s disordered domain substantially impaired CKAP2 localization to the axoneme. Collectively, these results demonstrate that CKAP2 localization to primary cilia requires the same 21- amino acids responsible for microtubule binding and stabilization.

### CKAP2 and CKAP2-Like mutually compensate for each other in the regulation of ciliary length and ciliogenesis

To study the impact of loss of CKAP2, CKAP2-Like or both, we used previously generated CKAP2 knockout RPE-1 cells and generated CKAP2-Like deficient RPE-1 cells by CRISPR/Cas9- genome editing targeting exons 4 and 5, respectively (Fig. S2*D*). We also knocked out CKAP2-Like in one of our CKAP2 KO clones and isolated two CKAP2/Like-double KO clones (dKO1 and dKO2) (Fig. S2*E*). The complete absence of protein expression was confirmed by sequencing (Fig. S3*A* and *B*), western blot (Fig. S3*C*) and immunofluorescence analyses (Fig. S4*A* and *B*) for all isolated KO clones. We thus compared the ability of CKAP2 KO, CKAP2-Like KO and CKAP2/Like dKO cells to properly assemble primary cilia under conditions of serum starvation. Immunostaining against the basal body marker CEP170 and acetylated α-tubulin for all KO cells revealed no significant alterations in basal body formation and distribution compared to wild-type cells (Fig. 4*A*). Interestingly, we also did not observe changes in the percentage of ciliated cells (Fig. 4*B*) in CKAP2 or CKAP2-Like KO cells. However, we observed a decrease in the overall ciliogenesis rate for both CKAP2/Like dKO clones, which ranged from 37% to 48% (dKO1 and dKO2, respectively), compared to 74% for CKAP2 KO1 control cells and 63% for wild-type cells (Fig. 4B).

**Figure 4.**
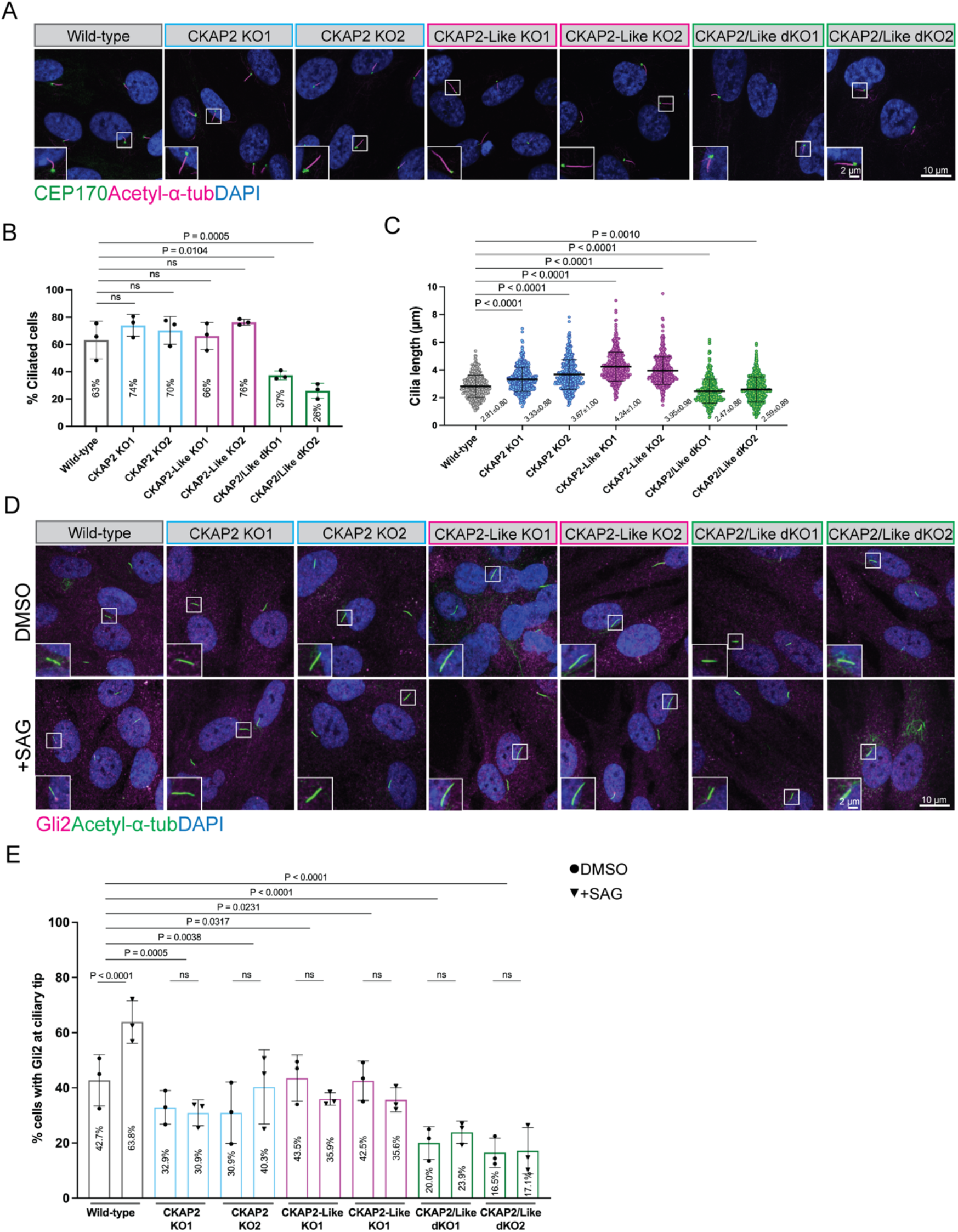
Knockout of CKAP2, CKAP2-Like or both promotes changes in ciliary length, ciliogenesis and signaling. **(A)** RPE-1 cells were serum-starved for 48 h, fixed and co-stained for endogenous CEP170 and acetylated-α-tubulin. **(B)** Quantification of the percentage of cells with cilia (WT n = 1509 cells; CKAP2 KO1 n = 1256 cells; CKAP2 KO2 n = 1454 cells; CKAP2-Like KO1 n = 1455 cells; CKAP2-Like KO2 n = 1310 cells; CKAP2/Like dKO1 n = 693 cells; CKAP2/Like dKO2 n = 751 cells). Data are representative of at least 3 independent experiments. **(C)** Quantification of ciliary length (WT n = 407 cells; CKAP2 KO1 n = 411 cells; CKAP2 KO2 n = 404 cells; CKAP2-Like KO1 n = 434 cells; CKAP2-Like KO2 n = 481 cells; CKAP2/Like dKO1 n = 451 cells; CKAP2/Like dKO2 n = 659 cells). Data are representative of at least 2 independent experiments. n.s. (not significant), P < 0.05 (statistically significant). **(D)** RPE-1 cells were serum-starved for 48 h and stimulated with DMSO (0.1%) or SAG (250 nM) for 1 h prior to fixing and co-staining for endogenous Gli2 and acetylated-α-tubulin. **(E)** Quantification of the percentage of cells with Gli2 at the ciliary tip in control (WT n = 458; CKAP2 KO1 n = 296; CKAP2 KO2 n = 406; CKAP2-Like KO1 n = 286; CKAP2-Like KO2 n = 431; CKAP2/Like dKO1 n = 284; CKAP2/Like dKO2 n = 249) versus SAG-treated cells (WT n = 392; CKAP2 KO1 n = 357; CKAP2 KO2 n = 375; CKAP2-Like KO1 n = 295; CKAP2-Like KO2 n = 319; CKAP2/Like dKO1 n = 258; CKAP2/Like dKO2 n = 226). Data are representative of at least 3 independent experiments. n.s. (not significant), P < 0.05 (statistically significant). Measurements are reported as average ± SD.

Since changes in ciliary length are a common feature of many ciliopathies (Deretic et al., 2023), we further measured the ciliary length of all KO clones (Fig. 4*C*). We found that CKAP2 and CKAP2-Like KO cells display an average increase in ciliary length of about 25.0% and 46.0%, respectively, compared to wild-type cells. To independently confirm the observed phenotypes, we performed acute knockdown of CKAP2 by siRNA, which resulted in unaltered ciliogenesis (Fig. S4*C* and *D*) and elongated cilia (Fig. S4*E*). This confirms that our CKAP2 KO phenotypes are not caused by CRISPR off-target effects. Overall, these data show that loss of either paralog consistently increases ciliary length, a surprising result given CKAP2/CKAP2-Like’s known roles in microtubule growth and stabilization (Lyalina et al., 2025; McAlear and Bechstedt, 2022; Paim et al., 2024).

To test the possibility that a mutual compensatory mechanism may drive the elongated cilia in cells lacking CKAP2 or CKAP2-Like, we measured the ciliary length for the remaining ciliated dKO cells and the presence of CKAP2 in the cilia of CKAP2-Like KO clones (and vice-versa). We found that dKO1 and dKO2 cells exhibit a ciliary length comparable to wild-type cells (Fig. 4*C*). Moreover, depletion of either MAP leads to upregulation of its counterpart within the ciliary compartment, as evidenced by a 38.5% increase in CKAP2-Like levels at primary cilia in the absence of CKAP2 (Fig. S5*A* and *B*). Likewise, we found a significant 26% increase in CKAP2 levels at the primary cilia in CKAP2-Like KO clones (Fig. S5*C* and *D*). Taken together, these data suggest that CKAP2 and CKAP2-like act as paralogs in stabilizing the axoneme, compensating for each other’s loss in maintaining ciliary length and ciliogenesis.

Next, we investigated whether the absence of CKAP2 or CKAP2-Like in primary cilia affects the translocation of Gli2, a canonical activator of the Hedgehog signaling pathway (Traiffort et al., 2010). Gli2 moves along the axoneme via intraflagellar transport (IFT) machinery and reaches the ciliary tip (Ku et al., 2025), where it is concentrated by the non-motile kinesin Kif7 (Blasius et al., 2021; Haque et al., 2026; He et al., 2014; Yue et al., 2022). We therefore performed an immunostaining analysis for Gli2 and acetylated α-tubulin in all KO cells. Activation of the Hedgehog pathway using SAG, a Smo agonist, did not significantly enrich Gli2 at the ciliary tip of CKAP2 KO, CKAP2-Like KO and CKAP2/Like dKO cells as compared to wild-type cells (Fig. 4*D* and *E*). These results suggest that loss of CKAP2, CKAP2-Like or both MAPs impairs the recruitment of Gli2 at the ciliary tip.

## Discussion

Here we show that CKAP2 and CKAP2-Like localize to primary and motile cilia of human cells (Fig 1*A* and *B*; Fig. S1*A* and *B*). To our knowledge, the presence of CKAP2 and CKAP2-Like as components of motile cilia and sperm flagella has not been previously reported (Blackburn et al., 2017; Broadhead et al., 2006; Cole et al., 1998). In the efferent ductules of human cells, both MAPs decorate the axoneme and the basal bodies of multiciliated (MCCs) and reabsorptive primary ciliated cells (RPCCs). We also observed CKAP2 and CKAP2-Like localizing to flagella in sperm cells navigating the duct lumen. In the epididymis, both MAPs are found in the primary cilia of principal and basal cells (Fig 1*B*; Fig. S1*B*). Consistent detection of CKAP2 and CKAP2-Like at the ciliary tip and transition zone may suggest occasional compartmentalization of both MAPs within the ciliary structure (Hansen et al., 2025; Saunders et al., 2025).

The presence of CKAP2 predominantly in cycling RPE-1 cells (Fig. 2*A* and *B*, and *E-G*) is in agreement with previous data reporting CKAP2-Like’s role in the cell division of the neural progenitor pool (Yumoto et al., 2013) and with its presence in the axoneme and basal body of proliferating cells (Ford et al., 2018; Gupta et al., 2015; Hansen et al., 2025; Mick et al., 2015). This versatility as a dual regulator of primary cilia and the mitotic spindle ensures stepwise control of the cell cycle progression (Jeon et al., 2006; Kwon et al., 2024; Tsuchihara et al., 2005) and faithful chromosomal segregation (Case et al., 2013; Hong et al., 2009; Paim et al., 2024; Yoo et al., 2016). Given that CKAP2 and CKAP2-Like are expressed throughout S/G2/M phases (Kwon et al., 2024; Paim et al., 2024), we propose that both MAPs may be involved in the ciliary dynamics after G1/S transition (Fig. 3*A* and *B*), temporarily stabilizing the axoneme and thus fine-tuning the timing of ciliary disassembly and mitotic entry (Ford et al., 2018). By acting as a microtubule polymerase (Lyalina et al., 2025), CKAP2 may counteract the runaway depolymerization of the axoneme during ciliary resorption. Following this process, CKAP2 populates the centrosomes, contributing to the assembly of the mitotic spindle shortly after cilia disassembly (Fig. 3*B* and Movie S2).

Structurally, CKAP2 and CKAP2-Like share 31% similarity and 13% identity along their protein sequences (Thompson et al., 1994). Interestingly, variabilities in phenotypic expression have been reported for patients with Filippi syndrome (Sabir et al., 2019; Sandhu et al., 2013; Yang and Marwaha, 2022) and also for CKAP2-Like knockout mice (Liu et al., 2026; Lyu et al., 2025), which may indicate a role for genetic compensation. Our results suggest that both MAPs have partially overlapping functions. CKAP2 compensates for the loss of CKAP2-Like (and vice-versa) in maintaining ciliary length and stability (Fig. 4*A-C*), which may be part of a safeguard mechanism that fine-tunes axoneme length. Indeed, it has been proposed that ciliary length is regulated by a precise balance between axoneme assembly and disassembly (Keeling et al., 2016). Several factors contribute to this regulatory process, with axonemal MAPs either controlling microtubule plus-end dynamics (Niwa et al., 2012; Saunders et al., 2025), competing for microtubule binding with other ciliary proteins or stabilizing the microtubule lattice (Ghossoub et al., 2013). We propose that CKAP2/CKAP2-Like may embody all these features while regulating axonemal microtubule dynamics by two distinct, but complementary mechanisms, as follows.

First, we found that cells lacking CKAP2 or CKAP2-Like develop elongated cilia (Fig. 4*A-C*), accompanied by increased ciliary enrichment of the remaining paralog (Fig. S5*A-D*). This suggests that in the absence of either paralog, its counterpart excessively promotes axoneme elongation (Fig. 4*A* and *C*), which may be achieved due to a combination of two factors: freeing of a CKAP2/CKAP2-Like binding site and local enrichment of the remaining paralog. The latter is supported by immunofluorescence results (Fig. S5*C* and *D*) and may be a consequence of an increase in transcriptional regulation (Kafri et al., 2005; Venkatesh et al., 2025), protein translation (Hao et al., 2021) or stability (Venkatesh et al., 2025), or recruitment of the non-ablated paralog to the ciliary compartment (Diss et al., 2014). By potentially catalyzing tubulin addition at the ciliary tip (Zhang et al., 2025) (Fig. 1*A* and Fig. S1*B*), both MAPs may ensure proper ciliary length together with other previously described MAPs (Saunders et al., 2025). In agreement with that, our recent findings demonstrate that CKAP2 targets the microtubule plus-end, where it promotes microtubule growth by stabilizing the transition state of an incoming tubulin dimer (Lyalina et al., 2025; McAlear and Bechstedt, 2022). This activity depends on a highly disordered domain of its structure, which we have shown here to be required for its ciliary localization (Fig. 3*C* and *D*).

Second, CKAP2 and CKAP2-Like distribute along the entire axoneme structure (b and c, Fig. 2*A*) and the combined loss of both paralogs suppresses ciliogenesis (Fig. 4*A* and *B*). This suggests that they may promote axoneme stabilization by directly interacting with and stabilizing the microtubule lattice (Zhang et al., 2025) (Fig. 2*A*; Fig. 3*A* and Movie S1). This is also supported by the observation that both MAPs are microtubule stabilizers when overexpressed in cells (Jin et al., 2004; Kwon et al., 2024; Yumoto et al., 2013).

The manner in which changes in ciliary length affect the Hedgehog signaling remains a matter of debate (Macarelli et al., 2023). We found that the elongation of cilia caused by the loss of CKAP2 or CKAP2-Like occurs alongside reduced accumulation of its canonical activator Gli2 at the ciliary tip (Fig. 4*D* and *E*). Given that Gli2 is primarily transported to the ciliary tip through interactions with IFT machinery (Ku et al., 2025), it is possible that the absence of each MAP induces specific changes in the microtubule state, thereby altering the binding or movement of signaling cargoes along the axoneme. This could affect the positioning of Gli2 at the ciliary tip, which would result in the dysregulation of Hedgehog signaling. In agreement with this, it has been recently shown that disrupting any paralog in mouse cells hinders the activation of this pathway by preventing the localization of Smo and Gli2 within primary cilia (Liu et al., 2026).

Overall, our results support a role for the MAPs CKAP2 and CKAP2-Like as potential axonemal stabilizers in primary cilia, in addition to their reported roles in the mitotic spindle (Kwon et al., 2024; Paim et al., 2024). Given the discrepancies in genotype-phenotype correlations associated with ciliopathies (Cardenas-Rodriguez et al., 2021), we propose that both MAPs precisely fine tune ciliogenesis, partially compensating for each other in human cells.

## Materials and Methods

### Cell Culture and Treatments

Human retinal pigment epithelial cells (hTERT RPE-1) were a gift from Arnold Hayer (McGill University) and were cultured in DMEM/F12 (11320082, Life Technologies) supplemented with 10% fetal bovine serum (FBS, 098-150, Wisent), 10 µg/mL hygromycin B and 1x penicillin-streptomycin (15140122, Life Technologies). All cell lines were maintained at 37 ^°^C and 5% CO_2_ in a humidified incubator (Thermo Fisher Scientific) and routinely trypsinized with 0.05% Trypsin-EDTA (25300054, Gibco). Ciliogenesis was induced by washing cells three times with DMEM/F12 without FBS. Cells were cultured for an additional 48 h in the absence of serum and fixed for immunofluorescence or harvested for western blot analysis. To assess Gli2 accumulation at the ciliary tip, cells were serum starved for 48 h and stimulated with DMSO (0.1%) or SAG (250 nM, 11914, Cayman Chemical) for 1 h.

### CRISPR/Cas9-genome editing

The knock-in of GFP to *CKAP2* by CRISPR/Cas9 has been previously described (Paim et al., 2024). For CRISPR/Cas9 knock-out, Cas9 Nuclease 2-NLS and sgRNAs targeting exon 4 of CKAP2 (sg1-GCAAAAAUAAUACAGUGGUG; sg2-GUCAAUUACUACAGUUGAAU; sg3- UCAACAUAUGACAUUAAGCC) or exon 5 of CKAP2-Like (sg1-UCAGGAAACAACUAGAAGAA; sg2-UUUGUUUUAAGUUCCAUAGG) were used. RNP complexes were formed in Nucleofector solution at a 6:1 ratio (sgRNA:Cas9) and immediately transfected by Nucleofection into RPE-1 cells using the P3 Primary Cell 4D-Nucleofector ® X Kit S (Lonza). EA-104 (RPE-1) program was set on the 4D-Nucleofector X unit (Lonza). RPE-1 cells were harvested, and 4 x 10^5^ cells were added to the RNP complex mix in a 16-well Nucleocuvette. The transfected mixes were resuspended in warm complete DMEM, plated in 12-well plates and media was changed 24h after nucleofection.

### Single-cell cloning

Stable knock-out cell lines were isolated from the edited pools by allowing cells to reach 70% confluency in a p100-dish. Single-cell cloning was then performed using the DispenCell Single-Cell Dispenser (Molecular Devices) according to the manufacturer’s instructions. Cells were individually seeded in a 96-multiwell plate containing conditioned media at 37 °C and the plate was monitored weekly for colony formation until 80% confluency was reached (around 2 weeks). Single-cell colonies were then expanded in 24-multiwell plates containing fresh media and further scaled up. After expansion, clones were sequenced and tested by western blot and immunofluorescence for the absence of CKAP2 and CKAP2-Like expression.

### Sequencing

To validate the genomic CRISPR edit presence, stable knockout cell lines were grown on p10- dishes for 48 h, and their DNA were collected with Monarch^®^ Spin gDNA Extraction Kit (T3010S, NEB). PCR was conducted with primers targeting a region on exon 4 for human CKAP2 (F- CTTCCTCCCCAAAAGGAAGCC; R-AAGCAGGCCTGGCTATAACTT) or on exon 5 of CKAP2- Like (F-GACACTGTTTATGGGCATGTCATT; R-GCCACGTGTGATCTTAGAATCTTG). PCR mixes containing 0.5 µL of Q5 polymerase (M0491S, New England Biolabs), 10 µL (1x Q5 reaction buffer), 1 µL (10 mM) dNTPs mix, 2 µL of genomic DNA, 2.5 µL (10 µM primers) and nuclease free H_2_O (up to 50 µL) were ran on a DNA Engine TETRAD2 Peltier Thermal cycler (MJ Research) with the following program: initial denaturation at 98°C for 30 s, 30 cycles of denaturation 98°C for 10 s followed by annealing at 67°C for 20 s and elongation at 72°C for 50 s, final elongation at 72°C for 3 min and a 4°C hold. PCR products were visualized on a 1 % agarose gel, purified using the Monarch^®^ Spin PCR & DNA Cleanup Kit (T1130S) and sent for Sanger Sequencing (Genome Quebec) with a nested sequencing primer for CKAP2 (CATGAGGCCCTTTCCGGATT) or CKAP2- Like (AAATTATAAGACTTTGGGCCAA).

### Immunofluorescence

Cells were cultured on glass coverslips and fixed with PFA 4% in PBS (J19943.K2, Thermo Fisher Scientific) for 10 min, washed twice with PBS and permeabilized with 0.5% Triton X-100 in PBS for 5 min at room temperature. Next, a blocking solution of 3% bovine serum albumin (BSA) (ALB001, BioShop) in PBS was added for 30 min at room temperature. Primary antibodies were diluted in blocking buffer and incubated for 2 h at 4 °C in a humidified chamber. Secondary antibodies were diluted in PBS and incubated for 1 h at room temperature in the dark. Nuclei were stained with DAPI (D21490, Invitrogen) diluted in PBS (1:1000) for 5 min. Glass coverslips were washed with PBS twice and mounted with FluorSave (345789, Millipore) onto glass slides. Primary antibodies diluted in blocking solution were: rabbit anti-CKAP2 (1:200, 25486-1-AP, Proteintech), rabbit anti- CKAP2-Like (1:200, 17143-1-AP, Proteintech), mouse anti-acetylated-α-tubulin (1:1000, sc-23950, Santa Cruz), rabbit anti-CEP170 (1:1000, 27325-1-AP, Proteintech), goat anti-Gli2 (1:150, AF3635, R&D Systems), mouse anti-Ki67 (#9449, 1:1000, Cell Signaling). Secondary antibodies diluted in PBS were: goat anti-mouse Alexa Fluor 488 (1:500, A11001, Invitrogen), goat anti-rabbit Alexa Fluor 647 (1:500, A21244, Invitrogen), donkey anti-goat Alexa Fluor 647 (1:500, A-21447, Invitrogen), donkey anti-mouse Alexa Fluor 568 (1:500, A-10037, Invitrogen).

### Immunohistochemistry

Human efferent duct and epididymal tissues were obtained from adult organ donors through the provincial organ donation program (Transplant Québec) under protocols approved by the Research Ethics Committee of the CHU de Québec–Université Laval Research Center (Approval No. 2018- 4043). Tissues from four donors (26, 32, 36, and 47 years of age) were included in this study. None of the donors had a documented history of reproductive disorders. Tissue procurement, microdissection, fixation, and formalin-fixed paraffin-embedding (FFPE) procedures were performed as previously described (Vinay et al., 2025). Briefly, formalin-fixed, FFPE sections (5– 10 µm thick) were deparaffinized in xylene, rehydrated through a graded ethanol series, and subjected to heat-induced antigen retrieval in 1× Tris-EDTA buffer (pH 9.0) at 110°C for 10 min. After cooling to room temperature, sections were permeabilized with 0.5% Triton X-100 in PBS for 15 min and blocked for 1 h in PBS containing 5% normal goat serum, 1% BSA, and Human TruStain FcX™ (1:100; 422302, BioLegend). Sections were incubated overnight at 4°C with primary antibodies diluted in blocking buffer as follows: mouse anti-acetylated α-tubulin IgG2b (1:2000, T7451, Sigma-Aldrich), mouse anti-γ-tubulin IgG1 (1:1000, ab27074, Abcam), rabbit anti-CKAP2 (1:200, 25486-1-AP, Proteintech), and rabbit anti-CKAP2-Like (1:200, 17143-1-AP, Proteintech). Following four 5-min washes in PBS, sections were incubated for 1 h at room temperature with secondary antibodies diluted 1:400 in blocking buffer: goat anti-mouse IgG1 Alexa Fluor 488 (A- 21121, Invitrogen), goat anti-mouse IgG2b Alexa Fluor 647 (115-607-187, Jackson ImmunoResearch), and goat anti-rabbit IgG Alexa Fluor 568 (A-11011, Invitrogen). Slides were mounted using VECTASHIELD Antifade Mounting Medium with DAPI (H-1200-10, Vector Laboratories).

### Live imaging of ciliary dynamics

To monitor primary cilia assembly and disassembly, RPE-1 cells were serum-starved for 32 h. CenSpark-650 was then added to culture media for 2 h at a final concentration of 100 nM prior to imaging. Cells were washed, and serum-starved medium supplemented with 25 nM of CenSpark- 650 was added for live imaging. Z-slices of 0.5 µm each were taken every 10 min for a total period of 20 h using a Nikon Eclipse Ti2 spinning disk confocal microscope (Cicero, Crest Optics) equipped with a 60x/1.40 NA objective lens and a stage top incubator (5% CO_2_ and 37 °C, UNO- T-H-PREMIXED, Okolab). Videos were processed using the Denoise.ai function of Nikon Elements AR 6.10.01 software and assembled using Fiji.

### Western Blot

Cells were cultured in 6-well plates until the desired confluency, quickly washed in cold PBS, and directly lysed in RIPA buffer (150 mM NaCl, 1.0% Triton X-100, 0.5% sodium deoxycholate, 0.1% SDS, 50 mM Tris, pH 8.0) containing protease inhibitor cocktail (P8340, Sigma-Aldrich). Cells were harvested with a cell scraper and lysates immediately centrifuged for 10 min at 4 °C and 14,000 rpm. Protein concentration was determined by Bradford using a BSA standard curve. A total amount of 20 µg of protein per sample was mixed with Laemmli loading buffer (200 mM Tris pH 6.8, 4% SDS, 40% glycerol, 4% 2-mercaptoethanol, 0.12 mg/mL bromophenol blue) and denatured for 5 min at 95°C. Proteins were separated on SDS-PAGE gel using a stacking (4% 29:1 acrylamide: Bis-acrylamide, 125 mM Tris pH 6.8, 0.1% SDS, 0.1% ammonium persulfate, 0.1% TEMED) and resolving gels (10% 29:1 acrylamide: Bis-acrylamide, 400 mM Tris pH 8.8, 0.1% SDS, 0.1% ammonium persulfate, 0.1% TEMED) at 120V. Transfer onto nitrocellulose membrane was performed for at least 16 h at 30V under wet conditions in 1X transfer buffer (14.4 g/L glycine, 3.0 g/L Tris, 20% methanol). The membranes were blocked with 5% nonfat dry milk in TBS-T 0.5% (50 mM Tris [pH 7.2], 150 mM NaCl, 0.5% Tween 20) for 2 h at room temperature. Three washes for 10 min each were performed after primary and secondary antibody incubation periods. Primary and secondary antibodies were diluted in blocking buffer and incubated for 1 h at room temperature. The following dilutions were employed: rabbit anti-CKAP2 (1:1000, 25486-1-AP, Proteintech), rabbit anti-CKAP2-Like (1:1000, 17143-1-AP, Proteintech), rabbit anti-TUBB3 (1:1000, 802001, Biolegend), and HRP anti-rabbit (1:10000, #1706515, Bio-Rad). The bands were finally visualized by enhanced chemiluminescence using Clarity (#1705060, Bio-Rad) and a Bio-Rad ChemiDoc MP.

### Molecular cloning

Full-length CKAP2 was obtained as described previously(McAlear and Bechstedt, 2022). CKAP2 domains were amplified from the full-length plasmid using Pfux7 polymerase and inserted into a pTWIST EF1alpha-Puro (Twist Bioscience) for mammalian expression as described previously(Lyalina et al., 2025). Correct inserts were confirmed by Sanger sequencing after bacterial transformation with plasmids and purification.

### Plasmid transfection

Plasmid DNA was transformed into NEB Turbo competent *E. coli* (C2984I, New England Biolabs), and minipreps were prepared using Presto™ Mini Plasmid Kit according to the manufacturer’s instructions. Plasmids used were CKAP2-mNG (aa 1-664), N-terminal α-helix CKAP2-mNG (aa 1- 60), disordered CKAP2-mNG (aa 61-345), Deleted SLiM CKAP2-mNG (aa 61-345, Δ169-189), α- helical bundle CKAP2-mNG (aa 346-549), and C-terminus CKAP2-mNG (aa 550-664). Transfection was performed in 8-well chambers (#94.6170.802, Sarstedt) containing 4.2 x 10^4^ cells per well using Xfect Transfection Reagent (631317, Takara Bio) with 300 ng of DNA following the manufacturer’s instructions. Cells were serum-starved 24 h post-transfection.

### RNAi-mediated knockdown

For transient knockdown, RPE-1 cells were cultured in complete medium for 24 h in a p6 or p12- multiwell plate containing glass coverslips. For siRNA transfection, 100-200 µL of Opti-MEM I (31985062, Gibco) and 4-8 µL of Lipofectamine RNAiMAX (13778150, Invitrogen) were added to tubes. siRNAs (60 pmol, stock solution 20 nmol) targeting CKAP2 mRNA (*Silencer* Pre-designed, AM16708, Thermo Fisher Scientific) (GGAUAGUAAUCAAACUCCGtt, CGGAGUUUGAUUACUAUCCtt), along with the Negative Control (Silencer No. 1, 440402, Thermo Fisher Scientific), were added to separate tubes and incubated for 10 min at room temperature. Media were changed shortly before transfection, and final mixtures were added dropwise. Cells were incubated for 24 h and serum-starved for an additional 48 h to induce ciliogenesis, as previously mentioned.

### Image acquisition, quantification and statistical analysis

Immunofluorescence tissue staining for CKAP2 and CKAP2-Like was assessed using a Zeiss LSM 900 confocal microscope equipped with an Airyscan 2 super-resolution detector. Images were taken using a 40x oil-immersion objective in Airyscan SR mode, with a pixel size of ≤0.04 µm. Z- stacking was carried out with optimized acquisition parameters to cover an approximate axial range of 10–20 µm. Images were subsequently processed and deconvolved using ZEN 3.2 Blue software (Carl Zeiss) and exported as TIFF files. Images were then used for figure preparation assembly. Primary cilia image acquisition was carried out using a Nikon Eclipse Ti2 spinning disk confocal microscope (Cicero, Crest Optics) equipped with a 60x/1.40 NA objective lens, AURA III Light Engine (Lumencor^®^) 549 and a Hamamatsu ORCA-Fusion C14440 camera with 110 nm pixel size controlled by Nikon Elements AR 6.10.01 software. The same number of confocal Z-stack slices (0.5 μm each) were acquired, and the maximum-intensity projection was used for fluorescence-intensity quantification. Images were acquired under the same gain and exposure time within the same experiment. All images were processed and analyzed using Fiji (ImageJ). Fluorescence intensity and cilia length were measured by using the segmented line tool (5-width). All statistical analyses and data plotting were performed using GraphPad Prism 10 (www.graphpad.com). In the dot plots, dots represent independent measurements grouped from at least two independent experiments. Mean and standard deviation were represented on graphs. For numerical data, Shapiro–Wilk normality tests were applied. For non-parametric data, Mann-Whitney U test was used where two independent groups were analyzed, while multiple comparisons were done using Kruskal-Wallis followed by Dunn’s Multiple Comparison test. For parametric data, One-Way ANOVA followed by Dunnett’s multiple comparison test was used when one independent variable was analyzed. Two-Way ANOVA followed by either Tukey’s or Šidák’s multiple comparison tests was used, when two independent variables were analyzed. A P-value less than 0.05 was considered statistically significant. All figures were assembled using Adobe Illustrator (v.29.4) and illustrations were created using BioRender.

## Acknowledgments

This work was funded by grants from the Canadian Institutes of Health Research (PJT-189995) and Natural Sciences and Engineering Research Council of Canada (NSERC; RGPIN-2024- 05603, RGPIN-2022-04551). We thank the members of the Bechstedt and Brouhard’s laboratories for constructive discussions on the project. The authors sincerely thank the nurses and surgeons at Transplant Québec for their essential contributions to tissue procurement and processing, and are deeply grateful to the donor families for their consent, which made this work possible. Finally, we would also like to thank Cédric Pourroy, Georgios N. Hatzopoulos, and Pierre Gönczy for kindly providing us with the CenSpark-650 probe.

## Author Contributions

LES: Data curation, formal analysis, investigation, methodology, validation, visualization, writing – original draft, writing – review and editing. GCS: Data curation, formal analysis, investigation, methodology, visualization. JA: Data curation, formal analysis, investigation. CB: investigation, methodology, visualization. SB: Conceptualization, funding acquisition, methodology, project administration, supervision, writing - original draft, writing – review and editing.

## Competing Interest Statement

The authors declare no competing interests.

## Data availability

The data underlying the graphs generated and/or analyzed during the current study are available from the corresponding author upon reasonable request.

## Classification

Cell Biology

## Supporting Information for

**Fig. S1.**
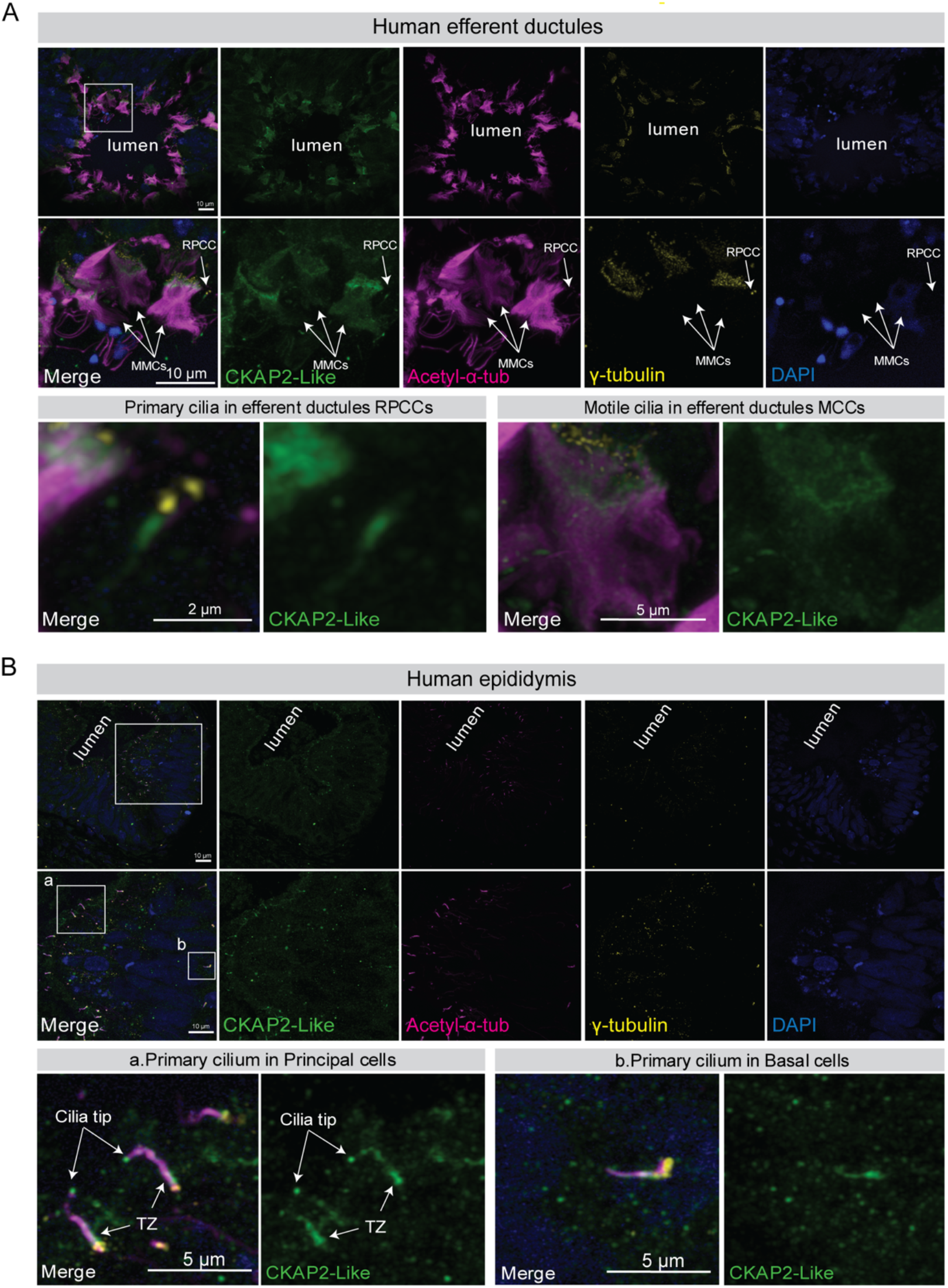
CKAP2-Like is found in the motile and primary cilia of the human male reproductive tract. **(A)** Confocal imaging microscopy of human efferent ductules stained for CKAP2-Like, acetylated-α-tubulin (axoneme) and γ-tubulin (basal bodies). Insets indicate CKAP2-Like localization in multiciliated cells (MMCs) and in reabsorptive primary ciliated cells (RPCC). Multiciliated cells are characterized by the apical alignment of numerous γ-tubulin-positive basal bodies. CKAP2-Like is also present in sperm flagella of the duct lumen. **(B)** Confocal imaging microscopy of human epididymis stained for CKAP2-Like, acetylated-α-tubulin (axoneme) and γ- tubulin (basal bodies). Insets indicate CKAP2-Like localization in the primary cilia of principal and basal cells. Occasional CKAP2-Like localization at the transition zone (TZ) and ciliary tip can be observed in principal cells.

**Fig. S2.**
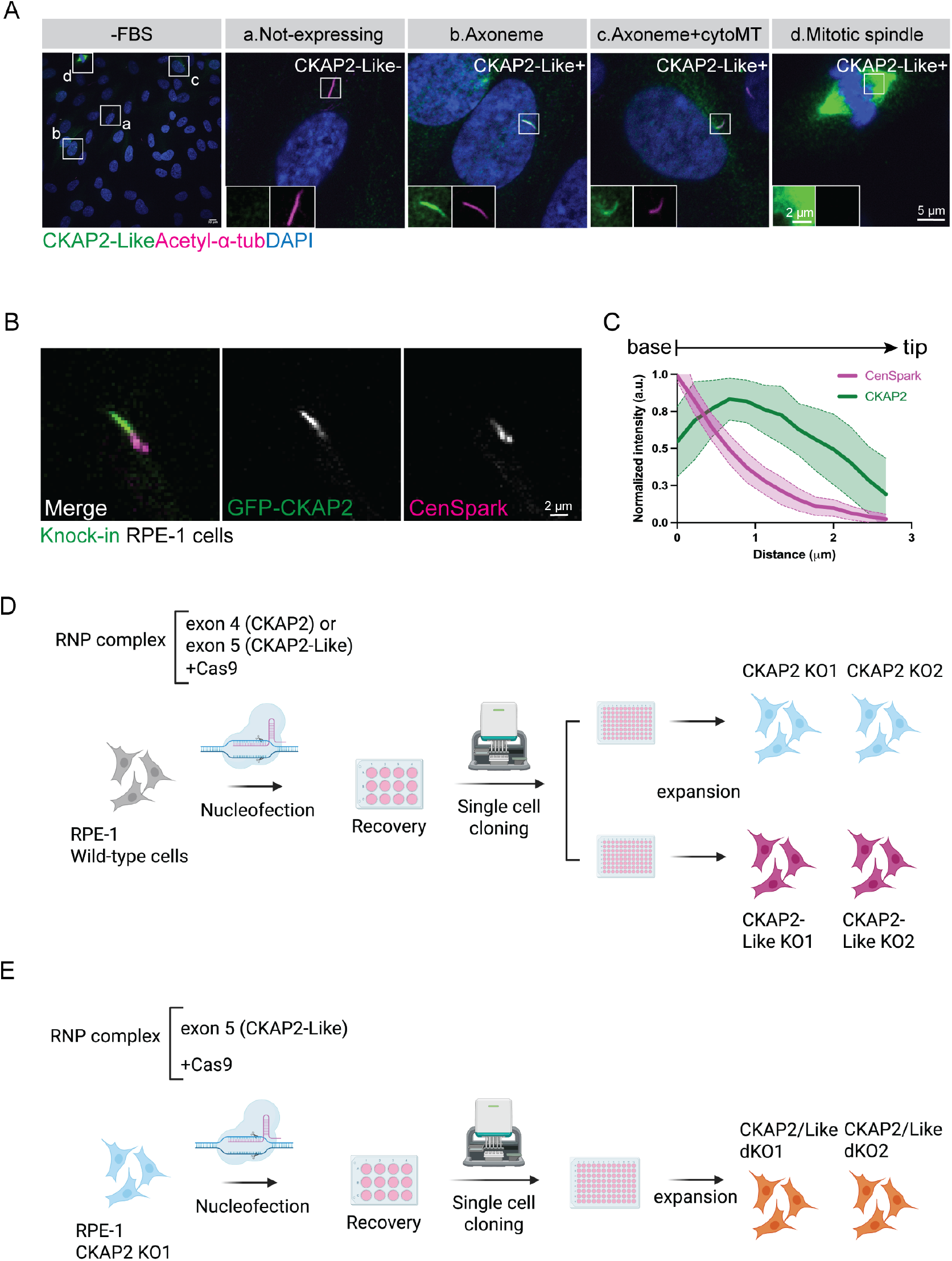
CKAP2-Like localization to the primary cilia is linked to cell cycle progression. **(A)** Wild-type RPE-1 cells were serum starved for 48 h (-FBS), fixed and co-stained for endogenous CKAP2-Like and acetylated-α-tubulin. Insets indicate different CKAP2-Like localization patterns. **(B)** GFP-CKAP2 knock-in RPE-1 cells were serum starved for 48 h and CenSpark-650 was added to media prior to live-imaging. **(C)** Normalized fluorescence intensity measured along the axoneme (from the ciliary base to tip) for n = 22 cells. **(D)** A Cas9 nuclease and sgRNA against the *CKAP2* and *CKAP2-Like* were transfected into RPE-1 cells by nucleofection. Single cell cloning was then performed into 96-well plates and knockout efficiency was confirmed in at least two clones by DNA sequencing immunofluorescence, and western blot. **(E)** A Cas9 nuclease and sgRNA against the *CKAP2-Like* were transfected into CKAP2 KO1 cells by nucleofection. Single cell cloning was then performed into 96-well plates and knockout efficiency was confirmed in at least two clones by DNA sequencing and western blot.

**Fig. S3.**
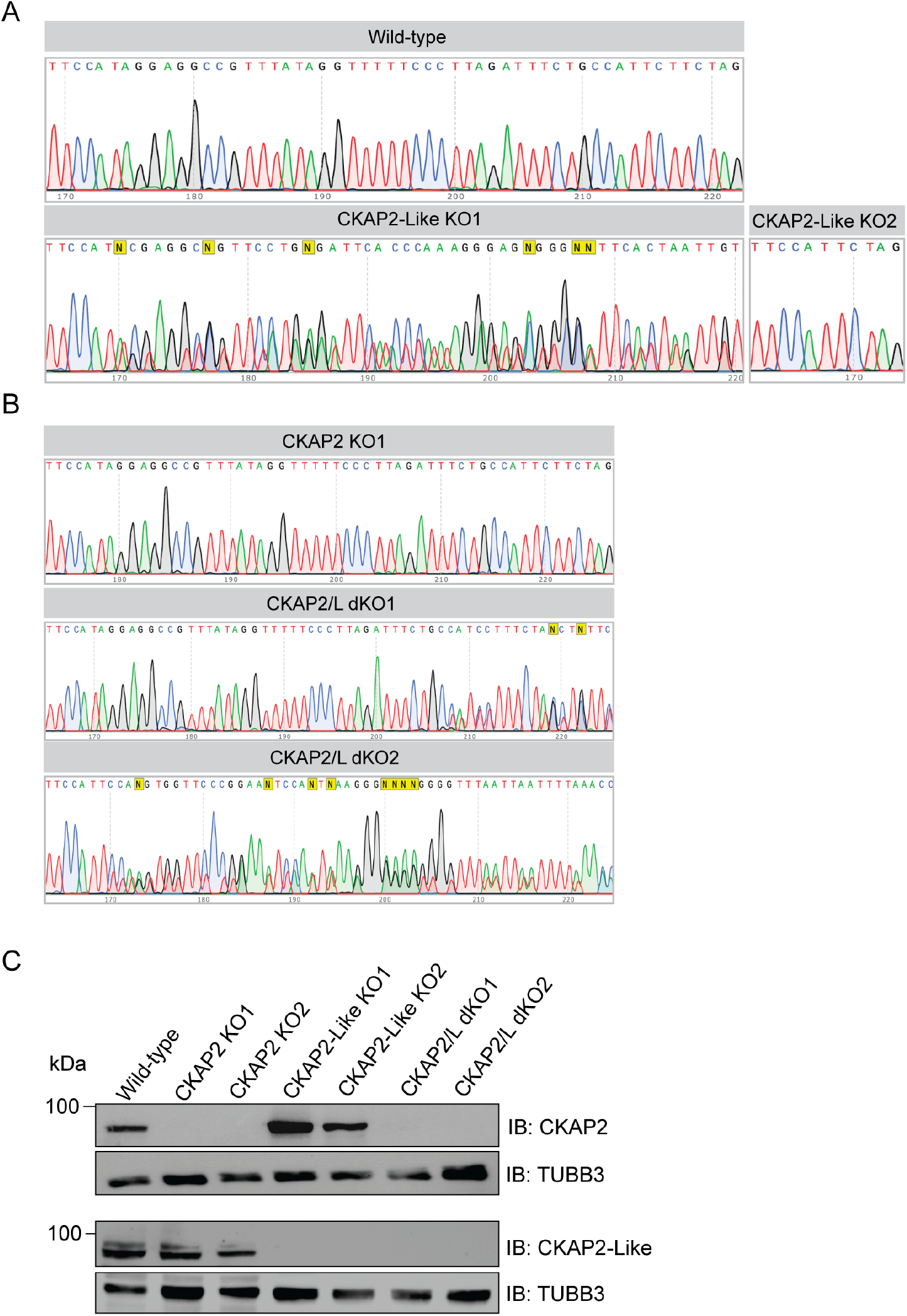
Validation of CKAP2, CKAP2-Like and CKAP2/CKAP2-Like double knockout clones. **(A)** Alignment results for the edited region corresponding to exon 5 in CKAP2-Like sequence of wild-type RPE-1 cells. **(B)** Alignment results for the edited region corresponding to exon 5 in CKAP2-Like sequence of CKAP2 KO1 clone. **(C)** Western blot analysis for CKAP2 and CKAP2- Like expression in WT, CKAP2 KO1, CKAP2 KO2, CKAP2-Like KO1, CKAP2-Like KO2, CKAP2/L dKO1 and CKAP2/L dKO2 cells. TUBB3 was used as a loading control.

**Fig. S4.**
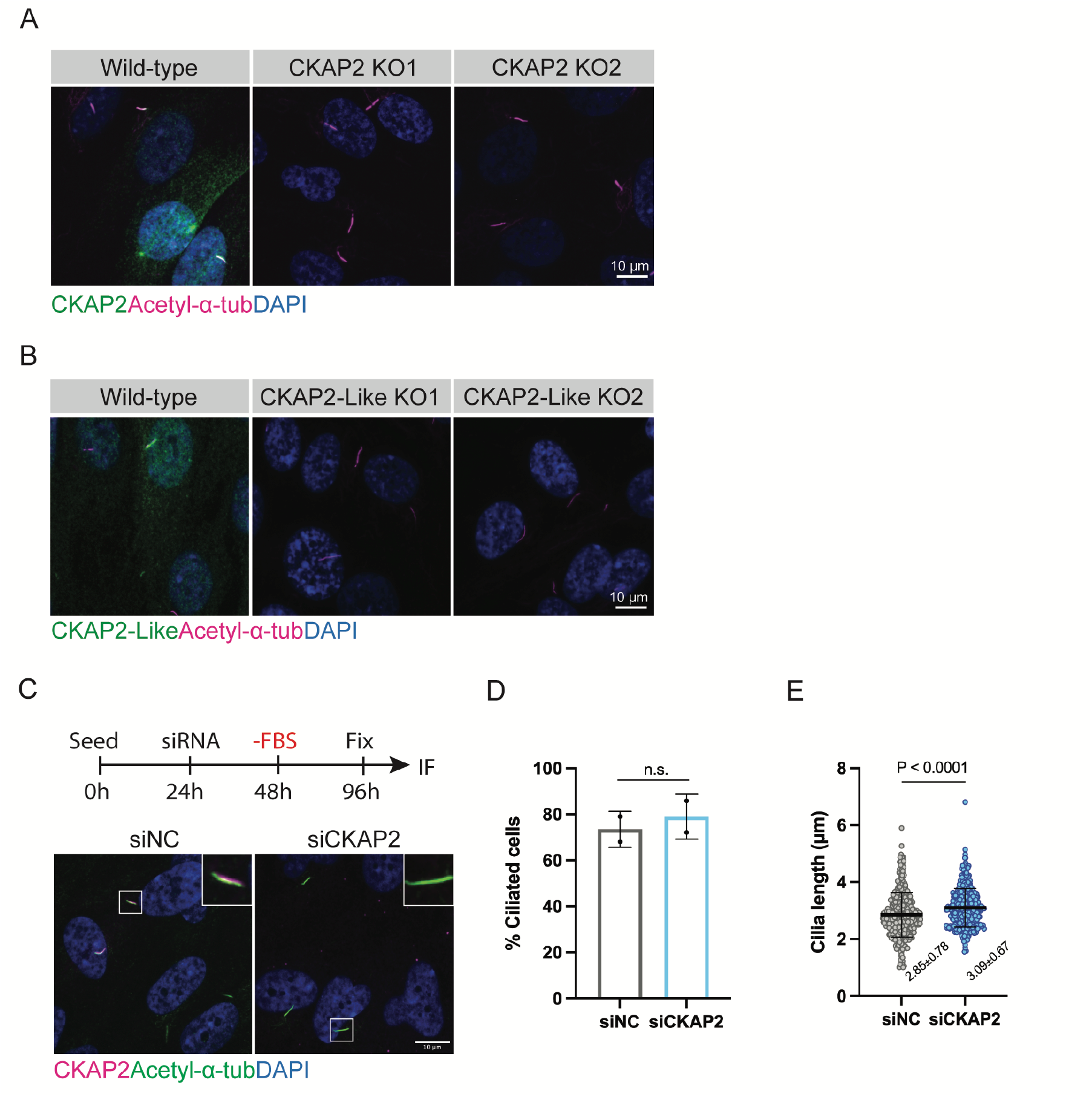
Knockdown of CKAP2 by RNAi leads to unaltered ciliogenesis and increased ciliary length. **(A)** Validation of CKAP2 expression levels at primary cilia in CKAP2 KO1 and CKAP2 KO2 clones by immunofluorescence. **(B)** Validation of CKAP2-Like expression levels at primary cilia in CKAP2-Like KO1 and CKAP2-Like KO2 clones by immunofluorescence. **(C)** Immunofluorescence analysis for CKAP2 and acetylated-α-tubulin in siNC and siCKAP2-treated cells. **(D)** Quantification of the percentage of cells with cilia (siNC n = 702 cells; siCKAP2 n = 768 cells). Data are representative of 2 independent experiments. **(E)** Quantification of ciliary length (siNC n = 379 cells; siCKAP2 n = 383 cells). Data are representative of 2 independent experiments. n.s. (not-significant), P < 0.05 (statistically significant). Nuclei were stained with DAPI

**Fig. S5.**
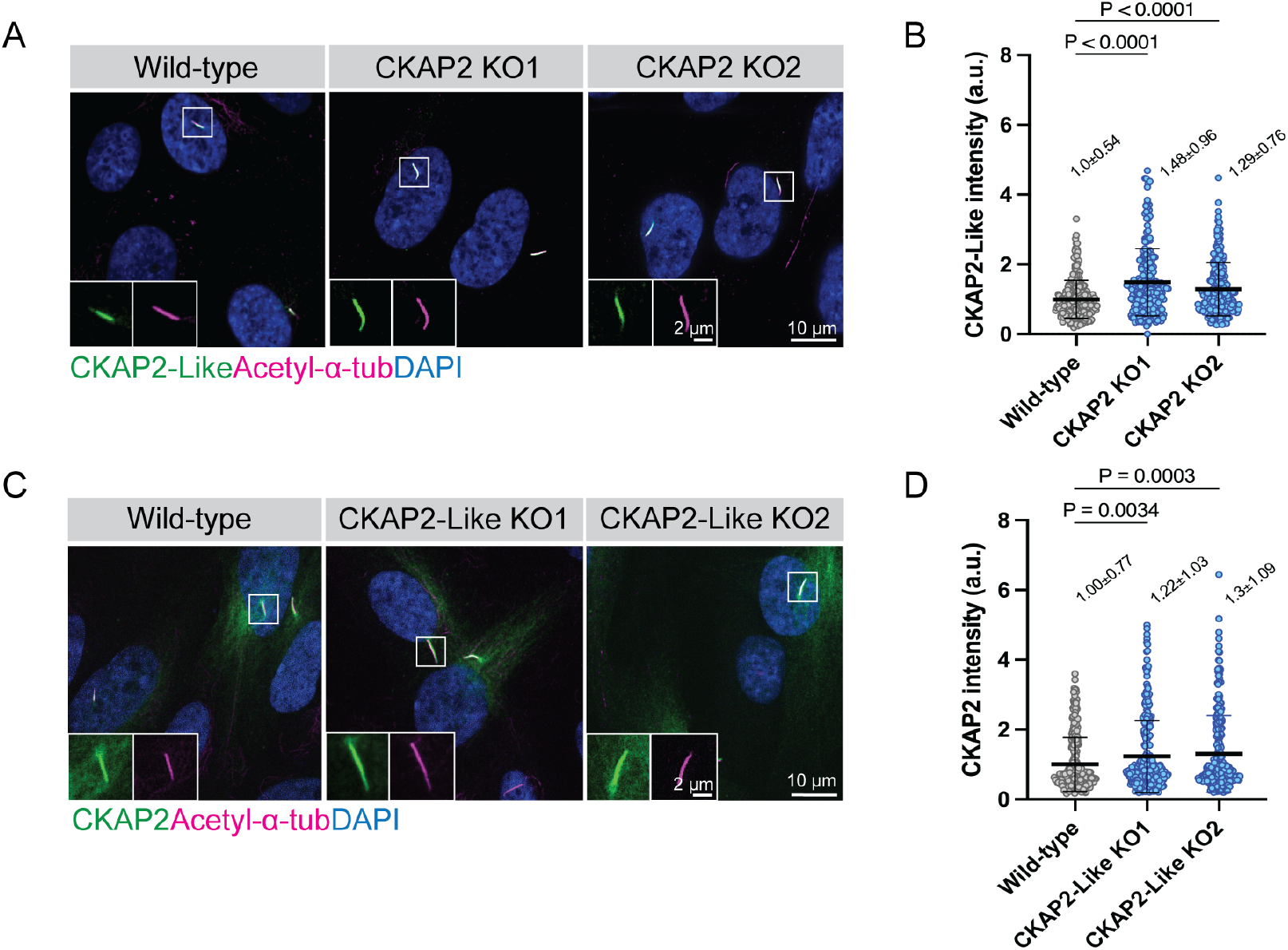
Knockout of either CKAP2 or CKAP2-Like leads to increased recruitment of its counterpart to the ciliary compartment. **(A)** WT and CKAP2 KO RPE-1 cells were serum starved for 48 h, fixed and co-stained for endogenous CKAP2-Like and acetylated-α-tubulin. **(B)** Quantification of CKAP2-Like fluorescence intensity at primary cilia of WT (n = 286 cells) and CKAP2 KO cells (KO1 n = 226 cells; KO2 n = 219 cells). Data are representative of at least 2 independent experiments. **(C)** WT and CKAP2-Like KO cells were serum starved for 48 h, fixed and co-stained for endogenous CKAP2 and acetylated-α-tubulin. **(D)** Quantification of CKAP2 fluorescence intensity at primary cilia of WT (n = 310 cells) and CKAP2-Like KO cells (KO1 n = 297 cells; KO2 n = 278 cells). Data are representative of at least 2 independent experiments. n.s. (not significant), P < 0.05 (statistically significant). Measurements are reported as average ± SD.

**Movie S1.** Dynamics of endogenously-labelled CKAP2 during ciliary assembly. Representative time-lapse video of GFP-CKAP2 knock-in RPE-1 cell, with CKAP2 undergoing ciliary localization after 32 h of serum starvation. CKAP2 initially localizes to the ciliary tip and then decorates the entire axoneme. CKAP2 = green (left inset), CenSpark = magenta (middle inset) and merge (right inset). Time is shown in minutes and video is accelerated to 1.5 frames per second.

**Movie S2.** Dynamics of endogenously-labelled CKAP2 during ciliary disassembly. Representative time-lapse video of GFP-CKAP2 knock-in RPE-1 cell, with CKAP2 localization during the transition between cilia disassembly and mitotic entry. CKAP2 follows the destabilization of the axoneme and populates the centrosomes. CKAP2 then fully decorates the mitotic spindle and later on chromatin, before being degraded at the end of mitosis. CKAP2 = green (left inset), CenSpark = magenta (middle inset) and merge (right inset). Time is shown in minutes and video is accelerated to 1.5 frames per second.

## Notes

### Competing Interest Statement

The authors have declared no competing interest.

